# PlantCV v4: Image analysis software for high-throughput plant phenotyping

**DOI:** 10.1101/2025.11.19.689271

**Authors:** Haley Schuhl, Keely E. Brown, Hudanyun Sheng, Parag K. Bhatt, Jorge Gutierrez, Dominik Schneider, Anna L. Casto, Lucia Acosta-Gamboa, Joe G. Ballenger, Fabio Barbero, Jackson Braley, Autumn M. Brown, Leonardo Chavez, Shannon Cunningham, Malinda Dilhara, Adam M. Dimech, Joseph G. Duenwald, Annika Fischer, Jared M. Gordon, Chloe Hendrikse, Gabriela L. Hernandez, John G. Hodge, Martina Huber, Brandon M. Hurr, Sanaz Jarolmasjed, Karina Medina Jimenez, Samuel Kenney, Grant Konkel, Alexander Kutschera, Sunita Lama, Matthew Lohbihler, Argelia Lorence, Collin Luebbert, Nathaniel Ly, Heather K. Manching, Annarita Marrano, Susan Meerdink, Nicholas M. Miklave, Pavan Mudrageda, Katherine M. Murphy, J. David Peery, Ronald Pierik, Seth Polydore, Caleb Robey, Tess Rogers, Tyler J. Schultz, Eliza Seigel, Dhiraj Srivastava, Stephan Summerer, Josh Sumner, Chong Teng, Adriane E. Thompson, Jose C. Tovar, Tim van Daalen, Mark Watson, John J. Wheeler, Mark C. Wilson, Kaitlyn R. Ying, Alina Zare, Yutai Zhou, Malia A. Gehan, Noah Fahlgren

**Affiliations:** Donald Danforth Plant Science Center, St. Louis, Missouri, USA; Compact Plants Phenomics Center, Washington State University, Pullman, Washington, USA; Maastricht University, Maastricht, Netherlands; Arkansas Biosciences Institute, Arkansas State University, Jonesboro, Arkansas, USA; University of Colorado, Boulder, Colorado, USA; Agriculture Victoria, Department of Energy, Environment and Climate Action, Bundoora, Victoria, Australia; Oklahoma State University, Stillwater, Oklahoma, USA; Plant Ecophysiology, Institute of Environmental Biology, Utrecht University, Utrecht, Netherlands; Syngenta Crop Protection AG, Basel, Switzerland; Collegiate School of Medicine and Bioscience, St. Louis, Missouri, USA; RAYN Growing Systems, Middleton, Wisconsin, USA; Independent; Purdue University, West Lafayette, Indiana, USA; North Carolina State University, Raleigh, North Carolina, USA; Department of Biology, University of Massachusetts, Boston, Massachusetts, USA; School of Earth, Environment, and Sustainability, University of Iowa, Iowa City, Iowa, USA; University of Louisiana at Lafayette, Lafayette, Louisiana, USA; University of Illinois, Urbana-Champaign, Illinois, USA; University of North Carolina, Chapel Hill, North Carolina, USA; University of Florida, Gainesville, Florida USA; ALSIA Centro Ricerche Metapontum Agrobios, Bernalda, Italy; Grinnell College, Grinnell, Iowa, USA; Wageningen University & Research, Wageningen, Netherlands; University of California, Davis, California, USA

## Abstract

PlantCV is an open-source Python project aimed at developing tools to address a range of image-based, plant phenotyping questions. PlantCV has been used for more than 10 years to automate trait collection from image data and the newest release, PlantCV version 4, continues to lower the barrier to entry for users without substantial coding experience through extensive example use-case tutorials and simplified installation. In addition to usability, we document added functionality since the release of PlantCV v2, including support for more image types such as fluorescence, thermal, and hyperspectral data. Finally, we describe the development of a new subpackage focused on morphological trait measurements like leaf angle, and demonstrate its utility as compared to more manual methods of data collection.

**CORE IDEAS:**

- PlantCV is an open-source, open-development, Python-based software package that has a new release for improved functionality and usability to make image analysis flexible and easier for researchers without a coding background.
- PlantCV is now capable of handling new data types that are relevant to researchers, such as thermal and hyperspectral, and has built in functionality for extracting information from these image types.
- The software project aims to lower the barrier to entry into image analysis for researchers by providing numerous, versioned, interactive tutorials that cover most common use cases, particularly in plant science.

## 1 INTRODUCTION

Phenotyping, especially high-throughput methods aimed at fast or easy approximation of traits or their proxies, has become a rapidly expanding area of plant science and is highly interdisciplinary (Pieruschka and Schurr, 2019). The field of phenomics focuses on the development of methods and tools to more accurately and efficiently describe an organism’s physical and physiological properties, including dynamically over time. Image analysis has gained traction as a phenotyping strategy because it has the potential to facilitate many trait measurements quickly and inexpensively (Li et al., 2020; Casto et al., 2021). Although image analysis approaches are often generalizable across domains, the target measurements are domain specific, and thus need methods specific to the plant science field (Li et al., 2020; Haase et al., 2022). Further, image analysis tools that have high flexibility like OpenCV (Bradski, 2000) and scikit-image (van der Walt et al., 2014) can be challenging for novice users with less data science expertise.

PlantCV is an open-source software package developed specifically for the extraction of quantitative trait data from images of plants (Fahlgren et al., 2015; Gehan et al., 2017). The vision of the PlantCV project is to develop phenotyping tools that are modular and reusable so they may be combined with ease to build flexible workflows to quickly extract biologically relevant data from images and sensors. In addition to providing sustained, functional tools, the PlantCV project aims to make phenotyping tools usable to researchers with broad backgrounds and varied expertise in data science and bioinformatics. The first version of PlantCV was released alongside the introduction of the Bellwether Phenotyping Facility in 2015 (Fahlgren et al., 2015), and has since supported a range of use cases, including image preprocessing for computer vision and machine learning applications (Agnew et al., 2016; Ubbens and Stavness, 2017; Liang et al., 2018; White et al., 2020; Nurminen and Malhi, 2020; Badhan et al., 2021; van de Koot et al., 2021; Kienbaum et al., 2021; Manss et al., 2023; Tanaka et al., 2023; Liu et al., 2023). The next major release, PlantCV v2.0 (Gehan et al., 2017), focused on developing modules that would increase the functionality of PlantCV, including image normalization (e.g., white balancing and color correction), auto-thresholding, size calibration using detectable reference markers, multi-plant detection and analysis, pseudo-landmarking for morphometrics, and a naive Bayes machine learning classifier (Gehan et al., 2017). The naive Bayes classifier included in PlantCV v2.0 (Gehan et al., 2017) has been used in biological studies to quantify plant damage by abiotic (Enders et al., 2019; Tovar et al., 2020; Ludwig et al., 2023; Ricono et al., 2025; Teng et al., 2025) and biotic stressors (Zheng et al., 2019). More recently, we developed the open-source R package pcvr, which enables users to perform statistical analysis on the quantitative data output from PlantCV workflows (Sumner et al., 2023).

PlantCV has successfully lowered the barrier to entry for extracting biologically meaningful traits from images of plants by providing flexible and modular functions that enable segmentation and measurement (Pierz et al., 2023; Lee et al., 2024; Teng et al., 2025; Alhassan and Aljahdali, 2025). However, the application of image analysis results from PlantCV has been limited by the types of traits that are possible to measure from RGB (red, green, blue) images. When studying stress tolerance, for example, differences in overall plant size, color, and leaf shape measurements between treatments may describe the response to an applied stress (Humplík et al., 2015; Nabwire et al., 2022; Ludwig et al., 2023; Duenwald et al., 2025), but may not provide information about the response mechanism. For drought stress specifically, researchers have supplemented these traits with thermal imaging (Chaerle et al., 2006; Granum et al., 2015) because estimated differences in leaf temperature can provide insight into stomatal conductance (Leinonen et al., 2006; Marchin et al., 2022).

Shifts in the color of leaf tissue from green to yellow can be symptomatic of several stresses including nutrient deficiency (Wiwart et al., 2009) and temperature stress (Enders et al., 2019; Ludwig et al., 2023). Imaging that includes reflectance in the near-infrared and red regions allows for the calculation of the normalized differential vegetation index (NDVI) as a more precise metric for general plant health without mechanistic information (Tucker, 1979). To determine if a shift in color corresponds to a change in a plant’s photosynthetic machinery, metrics like F_v_/F_m_ (a measure of the maximum potential of Photosystem II) and nonphotochemical quenching (NPQ, which describes excess light energy dissipated into heat) can be manually measured with chlorophyll fluorometry (Baker, 2008). Alternatively, chlorophyll fluorescence can be imaged using specialized camera systems (Awlia et al., 2016; Schneider et al., 2019; Casto et al., 2022).

Finally, precisely describing plant color might require more imaging than three wavelengths. Multi- or hyperspectral images containing information for up to hundreds of wavelengths have been used to calculate spectral indices that are predictive of biotic and abiotic stress and biochemical traits (Sarić et al., 2022). Machine learning classifiers using hyperspectral images have also been used to discriminate between roots of different species and live from dead roots (Baykalov et al., 2025). Because these images vary in the number of wavelengths discretely measured, using them for downstream analysis previously presented a challenge in PlantCV despite its flexibility.

We report here the next major release, PlantCV v4, which includes the ability to extract additional traits from RGB and grayscale data and functions to analyze new data types. For example, PlantCV was previously unable to easily measure morphological features like length, curvature, angle, and area of individual organs like leaves and stems, which is now possible through an added morphology subpackage. We have also added modules to support additional imaging modalities, such as thermal, fluorescence, and hyperspectral imaging. Application of these expanded imaging modalities for plant phenotyping is extensively reviewed elsewhere (Shakoor et al., 2015; Goggin et al., 2015; Mir et al., 2019; Casto et al., 2021).

A challenge with all software projects is sustainability, as many software projects, especially ones in academic research, are developed but not well documented, and then not maintained beyond publication (Lobet et al., 2013; List et al., 2017). Since its initial release on GitHub (see Data Availability) in 2014, PlantCV has been dedicated to developing infrastructure for long-term support and maintenance of the project (Fahlgren et al., 2015). In PlantCV v4, this goal has motivated the addition of stable releases that can be installed as packages from the Python Package Index (PyPI) (Python Software Foundation,) and conda-forge (conda-forge community, 2015), and the standardization of function input and output formats so that further improvements will remain compatible, which are described in this paper. We have limited breaking changes to the transition between major releases, which improves long-term stability of the codebase. Computational improvements in this version also speed up analysis of large datasets, enabling higher throughput. An improved set of tutorials, newly expanded phenotyping capabilities, and additional integration for software maintenance are included in this release and aim to grow PlantCV’s user base.

## 2 MATERIALS & METHODS

### Plant growth and imaging

Rice image data was previously published, including detailed methods (Huber et al., 2024). Images from plants 14 days after sowing were used in this manuscript to demonstrate the utility of the PlantCV v4 morphology subpackage.

### Color correction

Image color correction was done using Macbeth ColorChecker cards (McCamy et al., 1976). PlantCV v4 can automatically detect color cards using the standard MacBeth ColorChecker layout of 24 color chips in a 4 x 6 grid layout. Specific color card models tested were ColorChecker Classic (Calibrite LLC, formerly X-Rite, Inc.), ColorChecker Passport Photo 2 (Calibrite LLC, formerly X-Rite, Inc.), ColorChecker Classic Nano (Calibrite LLC, formerly X-Rite, Inc.), ColorChecker Classic Mini (Calibrite LLC, formerly X-Rite, Inc.), and 24ColorCard-2×3 (CameraTrax).

### Ground-truth leaf insertion angle estimation with ImageJ

We collected manual digital measurements for rice leaf insertion angle using ImageJ (Schneider et al., 2012) for the images from Huber et al., 2024. Using the angle tool, we clicked on the culm of the plant at the point where the focal leaf connects to the culm and a point on the leaf where the blade had not begun to bend to capture the insertion angle and not a measure of leaf erectness (Figure S1). We repeated this process on both the 2nd and 3rd leaves from the top when present. In total, we measured 46 images with 2 plants per image and 1 to 2 leaves per plant, totaling 139 leaf measurements.

### Morphology Measurements with PlantCV

Starting with images of rice plants, we converted the images to grayscale using the “cyan” color channel from the CMYK color space and then thresholded on pixel intensity to segment plants from the background. Using regions of interest (ROIs), we further separated plants from pots, stakes, and other background features. Once a cleaned binary mask of the segmented plant was extracted, a skeletonization method was used to thin the mask to a single pixel thickness. Since skeletonizing an image involves making iterative passes to remove the borders until only the structure of the object is remaining, any boundary noise (jagged edges) in a plant mask will result in spurious branches on the skeleton, which we pruned away using a branch size threshold of 100. We then sorted skeleton segments into leaves and culms. Finally, we extracted leaf insertion angle by calculating the angle between the culm and the first 100 pixels of each leaf segment. Workflows used to analyze rice image data are available on GitHub (see Data Availability).

### Hyperspectral Image Data

PlantCV was tested with multi and hyperspectral image data from a range of camera sensors, including Headwall Scientific (VNIR E-Series and SWIR), Teledyne (PyLoN series), Specim (IQ), Fotenix (Delta MkII), Cubert (Ultris S5), and RAYN Growing Systems (RAYN Vision System). Hyperspectral data are stored in binary interleaved formats, and PlantCV is compatible with the three common types: Band Interleaved by Line (BIL), Band Interleaved by Pixel (BIP), and Band Sequential (BSQ). PlantCV reads hyperspectral raster data and metadata in the Environment for Visual Imagery (ENVI) and Esri ASCII formats.

## 3 RESULTS AND DISCUSSION

Changes made to the infrastructure of PlantCV v4 from PlantCV v2 improved software sustainability, including exclusive compatibility with Python 3 consistent with the deprecation of Python 2, improved unit testing coverage, simplified installation, and improved documentation. Development of PlantCV includes community contributions and uses automatic quality control methods through GitHub Actions. The requirement for documentation of all new contributions makes it straightforward to track progress toward stable releases (Figure 1). Users of PlantCV can expect support in the form of documentation on the use of every function, a detailed changelog for each released version, links to resources for getting help and contributing, tutorials, and recorded presentations and workshops. Here, we describe specific improvements to the software that expand the types of research that can benefit from using PlantCV, as well as expansion of data types compatible with existing workflow strategies.

**Figure 1.**
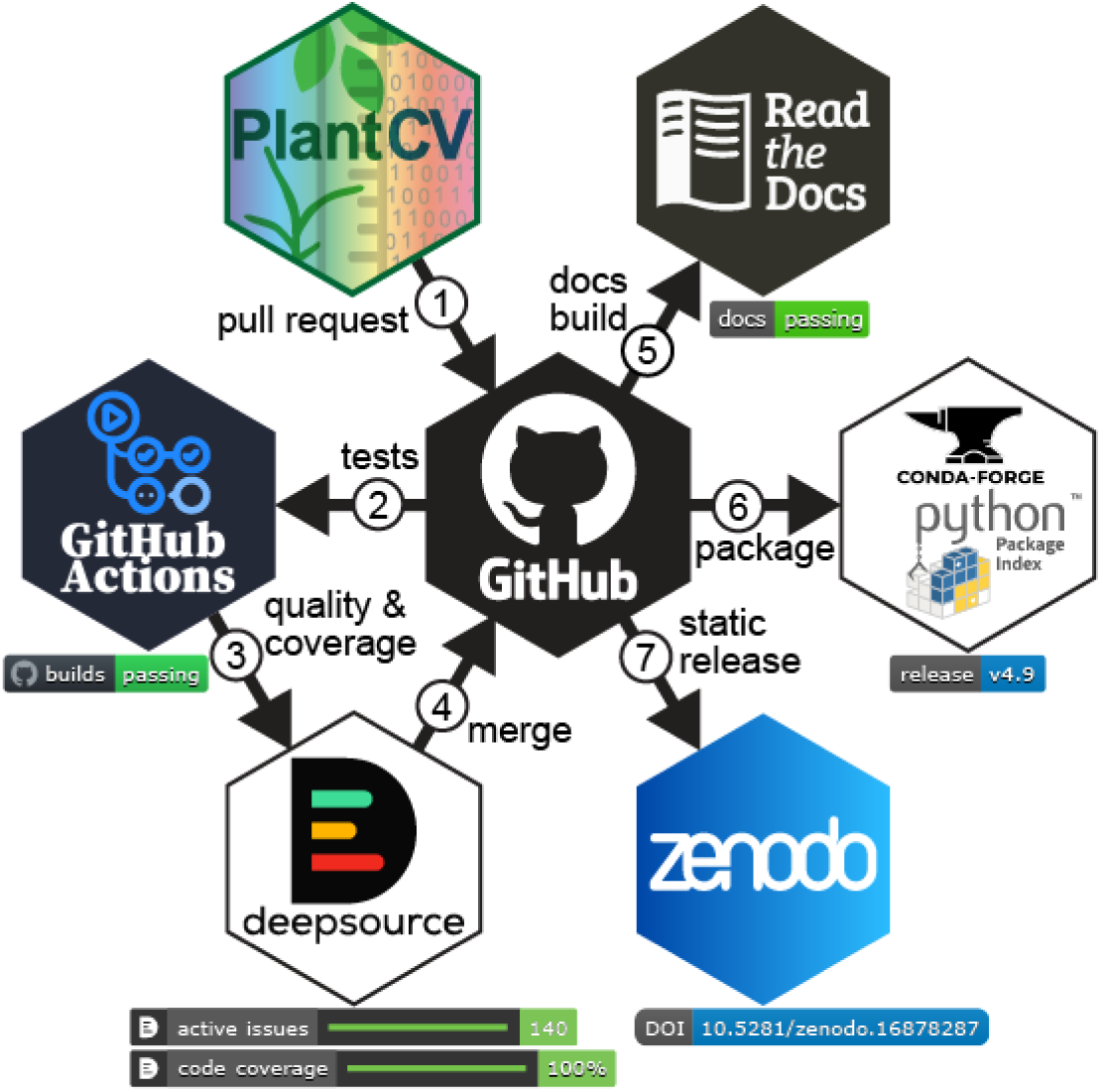
PlantCV development workflow. PlantCV development follows steps 1-7, resulting in a new version release.

### 3.1 Improved usability

#### Software installation and maintenance

Software that cannot be installed reliably and easily misses opportunities for use. PlantCV v2 had well-documented instructions for installation; in PlantCV v4 we added the routine release of stable packages that can be installed, along with automatic installation of dependencies from PyPI or conda-forge (Figure 1). By providing stable, cross-platform releases, installation is streamlined and users only need to follow the PlantCV instructions, rather than also needing to learn how to install all the dependencies separately.

#### Input consistency

PlantCV is a collection of modular functions that do discrete operations that can be linked together in custom arrangements to form a workflow (Figure 2). A PlantCV workflow is often developed in a Jupyter notebook environment by utilizing the outputs of one or more prior functions as the inputs for the next function in a sequence (Figure 2). To make PlantCV v4 more interoperable with other packages and internally consistent, we standardized the vocabulary of function input keywords to provide users immediate indications about the type of input data expected. For example, the keywords ‘img,’ ‘rgb_img,’ ‘gray_img,’ and ‘bin_img’ are used to indicate that a function can utilize multiple image types, only RGB images, only grayscale images, or only a binary image, respectively. In addition to the online documentation, each function now has built-in documentation (Python “docstrings”) that describes the inputs and outputs. PlantCV function docstrings can be accessed using the Python “help” function or using tooltips and autocompletion in the Jupyter environment (Kluyver et al., 2016) making it easier to quickly access documentation while building a workflow.

**Figure 2.**
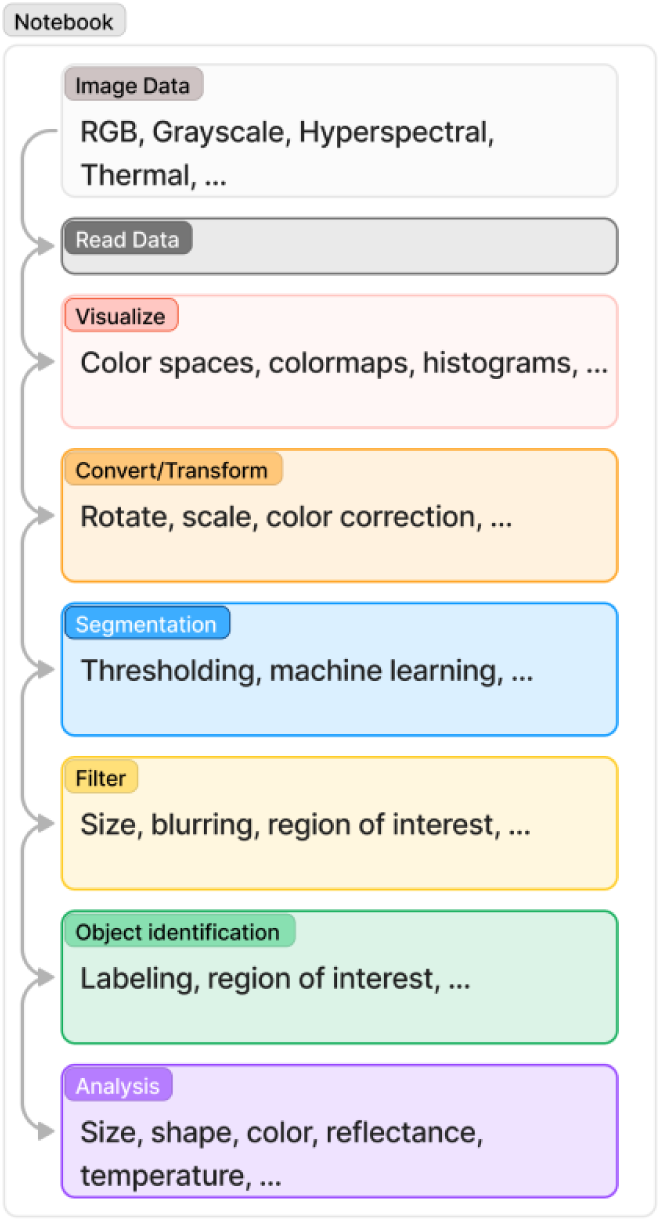
Generalized PlantCV workflow. PlantCV workflows are developed in Jupyter notebooks.

#### Output automation

In PlantCV v2, analysis functions returned lists of observations (numerical measurements or qualitative data) to the user, and the user was required to write code to save the results to a temporary file prior to loading into a SQLite relational database (Gehan et al., 2017). The database schema for the results tracking system was rigid and had to be updated whenever new measurement capabilities were added to PlantCV, and the method for saving results was not user-friendly. In PlantCV v4, all analysis functions that collect observations now utilize a background outputs recorder framework that collates workflow results automatically. The outputs framework stores observations using an extended trait definition schema that maps to attributes in the Minimum Information About a Plant Phenotyping Experiment (MIAPPE) vocabulary (Ćwiek-Kupczyńska et al., 2016). Each observation is recorded with the trait, method, scale, data type, value, and data labels. The PlantCV v4 outputs framework also provides a method for users to add custom measurements that are not built into PlantCV. This improved workflow flexibility is critical for the interoperability of PlantCV with other software so that users are not limited to recording observations from defined internal functions. The required metadata is designed to encourage interoperability, reproducibility, and interpretability of results.

#### Visualization for building workflows

PlantCV workflows are built from modular functions that typically require parameterization to segment focal objects from the background, filter out noise, and identify and measure properties of individual objects (Figure 2). To streamline the process of choosing effective parameters, visualization functions were added in PlantCV v4 to allow users to see and make data-driven decisions about parameter combinations. For example, workflows that use a thresholding strategy for image segmentation need to select one or more grayscale images derived from the input image color properties. In PlantCV v4, all available colorspace channels can be visualized in a single plot to observe an RGB image converted to HSV (hue, saturation, value), LAB (lightness, red-green, blue-yellow), and CMYK (cyan, magenta, yellow, black) color spaces, allowing users to identify specific channels that have the best contrast between the object of interest and the background (Figure 3A). Similarly, to better select a cutoff value for binary thresholding, a histogram visualization function was added that plots the pixel intensity values of a grayscale or RGB image (Figure 3B). Histogram visualization is also available for hyperspectral images when selecting individual bands. Other functions for improved visualization include displaying and charting object sizes, colorizing grayscale and binary images, and overlaying images.

**Figure 3.**
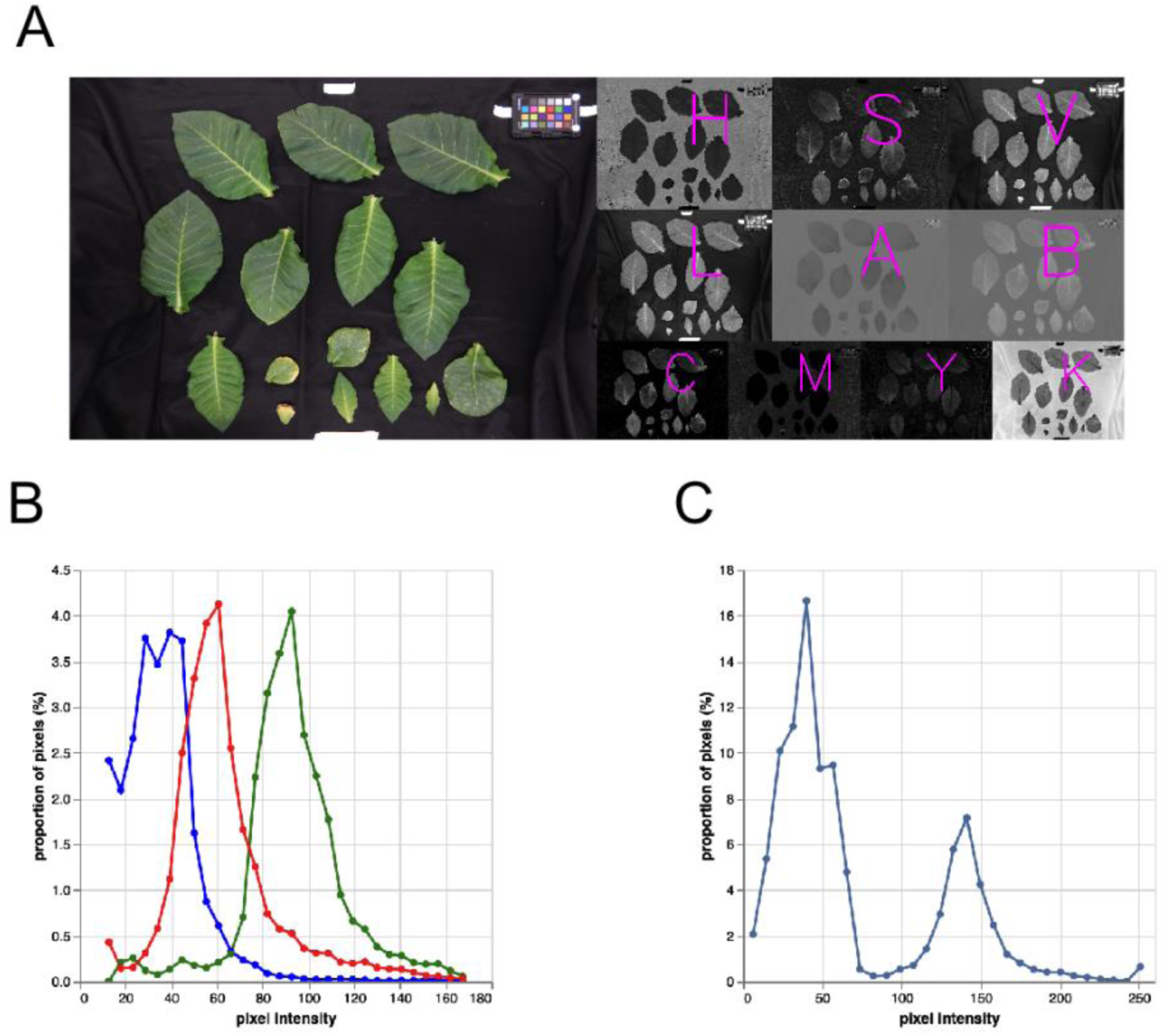
New visualization functions are useful for workflow development. (A) Colorspaces can be visualized simultaneously so that users can choose which grayscale conversion makes the most sense for segmenting focal objects from any background. (B, C) Plotted histograms of pixel values across RGB (B) and grayscale (C) color channels, as well as grayscale, can help users choose a threshold for segmentation, with the gray peak around 45 belonging to background and the peak around 140 belonging to plant material. In (B), red, green, and blue color channels are shown with matching colored lines. Image from (Murphy et al., 2025).

#### Computational improvements

The PlantCV v2 parallelization framework was limited to computing on a single multi-core/multi-CPU (central processing unit) machine. In PlantCV v4, we added a new framework based on the Dask package (Dask Development Team, 2016), which creates a cluster of worker processes that are used by the scheduler to batch process images. Now, analysis throughput can be improved by distributing workloads across multiple machines in a cluster managed by a supported distributed resource management system, which includes many commonly used job schedulers and workload managers. During parallelization, PlantCV automatically collects metadata, filters data, specifies computing requirements, processes images, and saves output images and measurements. Previously, these processes were parameterized using lengthy command-line options, which we replaced in PlantCV v4 by a configuration file. Now, a single line of code is used to run the workflow using the configuration file. Because this file is a record of all the parameters used during parallelization, it both eases use and helps to increase experimental reproducibility. Configuration files are also more approachable for users less familiar with the command-line and can be edited with any standard text editor.

### 3.2 Tools to increase the flexibility of workflows

#### Color correction

Variation in imaging or the imaging environment can result in image-to-image variability, which can impact downstream image processing. For example, we previously showed that variation in fluorescent light fixture output due to changes in ambient air temperature creates variation in image brightness (Berry et al., 2018). To standardize image color profiles across a dataset within PlantCV v4, we added a color correction module (Berry et al., 2018). Datasets that contain a reference color card in each image can be corrected using reference color values or a reference image, which was shown to significantly improve automatic thresholding methods and increase the accuracy of measured phenotypes, particularly hue (Berry et al., 2018). The individual color chips on a color card need to be identified and labeled in each image to extract the observed color values, which can be challenging if the position of a color card within an image is inconsistent. To address this issue, we added an algorithm to automatically detect color cards in PlantCV v4. The color card detector uses edge detection, and object shape and size filtering to identify the outlines of most of the individual color chips. Because the detector is not guaranteed to detect all the chips, the position and spacing of chips is estimated based on a 4×6 grid layout that is common to Macbeth ColorChecker cards (McCamy et al., 1976). The grid layout is then transformed, a combination of being rotated and resized using a perspective transformation method. This transformed grid is applied to match the outer boundaries of the detected chips within the image. The orientation of color card chip masks is inferred by which detected chip is most likely the white chip based on detected color values. Additionally, the height and width of the chips are estimated by recording the mean and median values observed, and the chip size can be used for normalization of other measurements, such as size conversion from pixels to cm, to the same scale for a dataset. We have successfully detected a variety of sizes of color cards of the Macbeth ColorChecker layout (McCamy et al., 1976) in variable imaging conditions (Figure 4). PlantCV color correction methods have also been used for applications outside of the plant science field (Paradis et al., 2020; Kwon et al., 2022).

**Figure 4.**
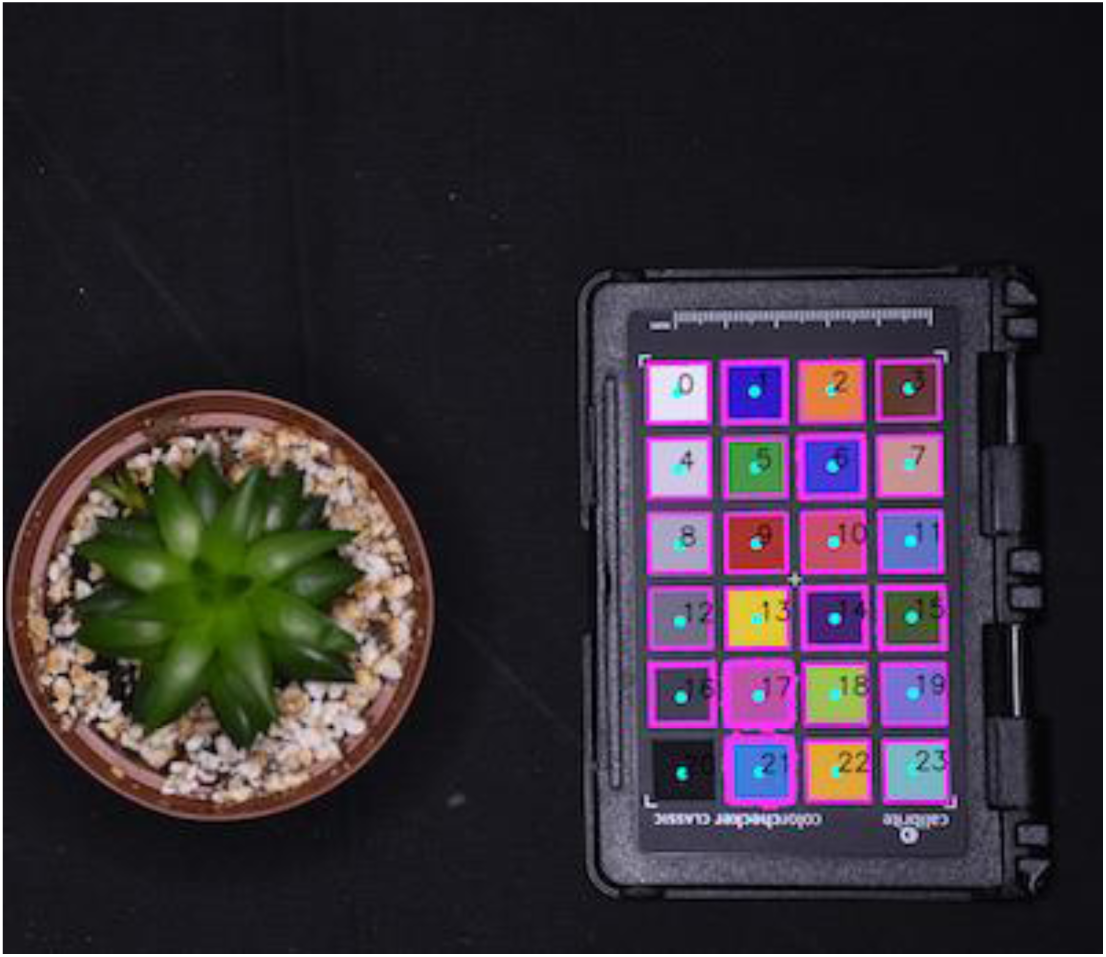
Color card detection for color correction. Pink outlines represent the detected objects in the image that the algorithm detects as color card chips. Cyan circles represent the area sampled for pixel values, and the color card chips are numbered. The orientation will not impact the order of cards detected with the latest version of this algorithm, since the white chip is identified from sampled pixels.

#### New Region of Interest (ROI) methods

After an object is segmented from the background, a “region of interest” (ROI) serves to spatially distinguish parts of an image for further filtering and measurement. Prior to v4, options for specifying ROIs were limited to circles, ellipses, and rectangles. In PlantCV v4, we added a custom ROI method that allows users to define a custom polygon by providing a list of coordinates around an object. Increased flexibility in target object selection can simplify workflows (Figure 5A).

**Figure 5.**
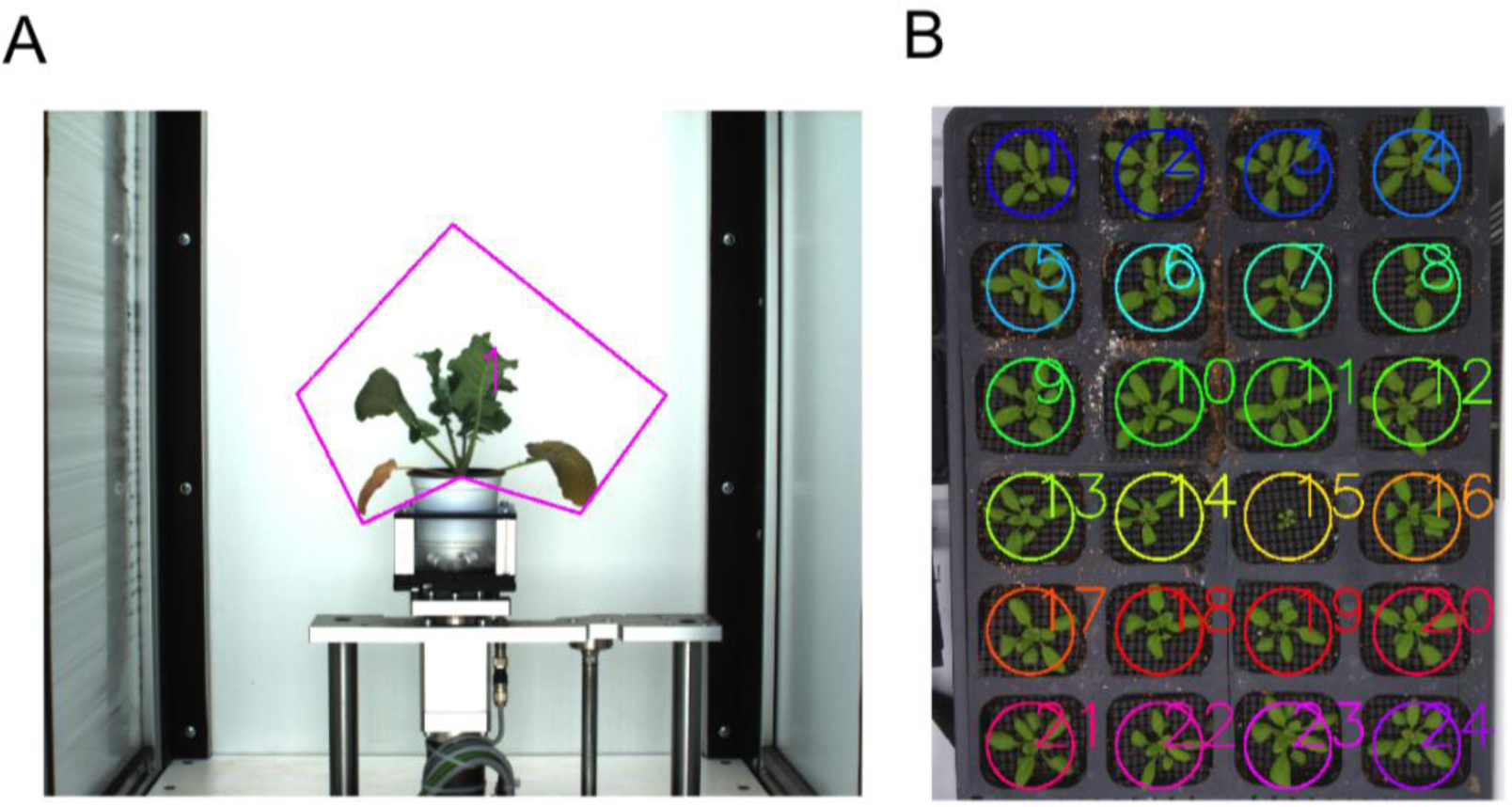
New ROI tools make both single and multi-plant analysis more flexible. (A) A custom ROI can be specified with as many vertex points as is required to enclose a focal object. Image from (Ricono et al., 2025). (B) A grid of circular ROIs can be easily created by specifying the number of rows and columns in an image. Image from (Panda et al., 2023).

In previous versions of PlantCV, analysis of multiple objects within an image required coding custom loops to iterate over a set of ROIs. In PlantCV v4, multiple ROIs can be used to create a labeled mask that uniquely identifies each object. All analysis modules that measure size, shape, color, spectra, etc. were refactored to accept labeled masks and automatically measure multiple individual objects without additional user coding.

In controlled plant growth environments, grid based experimental layouts are common, such as using a tray of plants or well plates, that warrant multiple ROIs in a specific grid format. In PlantCV v4, we streamlined the process of defining multiple regions of interest with tools that automatically define a grid of ROIs (Schuhl et al., 2022). Automated ROI tools create a grid of multiple circular or rectangular ROIs based on detection of the grid layout from a binary mask of objects and given the number of rows and columns provided by the user (Figure 5B). ROIs can also be created by detecting the circular wells in a laboratory sample plate.

### 3.3 Expansion of input data types

#### Fluorescence imaging for photosynthetic traits

Plant fitness traits, including measures like yield and abiotic stress, are well correlated with photosynthetic indices calculated from fluorescence imaging (Chaerle and Van Der Straeten, 2001; Kolber et al., 2005; Chaerle et al., 2006; Baker, 2008). Although previous versions of PlantCV included tools to analyze fluorescence imaging data, PlantCV v4 simplifies the process of combining the multiple image frames needed to calculate photosynthetic measurements such as F_v_/F_m_ (Baker and Rosenqvist, 2004). PlantCV now reads the raw binary image files and metadata from PhenoVation B.V. instruments and combines the chlorophyll fluorescence frames into a time series datacube. Updated analysis functions utilize the multi-image datacubes to calculate F_v_/F_m_, non-photochemical quenching (NPQ), and spectral indices like chlorophyll index and anthocyanin index (Casto et al., 2022; Murphy et al., 2025) (Figure 6A). The photosynthesis subpackage records traits including the mean, median, mode, and maximum F_v_/F_m_ and NPQ values for objects of interest in the image, as well as full histograms for each measurement. The function also prints out a debugging image with the histogram data that can be optionally saved. Tutorials on analysis of photosynthetic measurements are available (annikafischer et al., 2025; Schuhl et al., 2025a).

**Figure 6.**
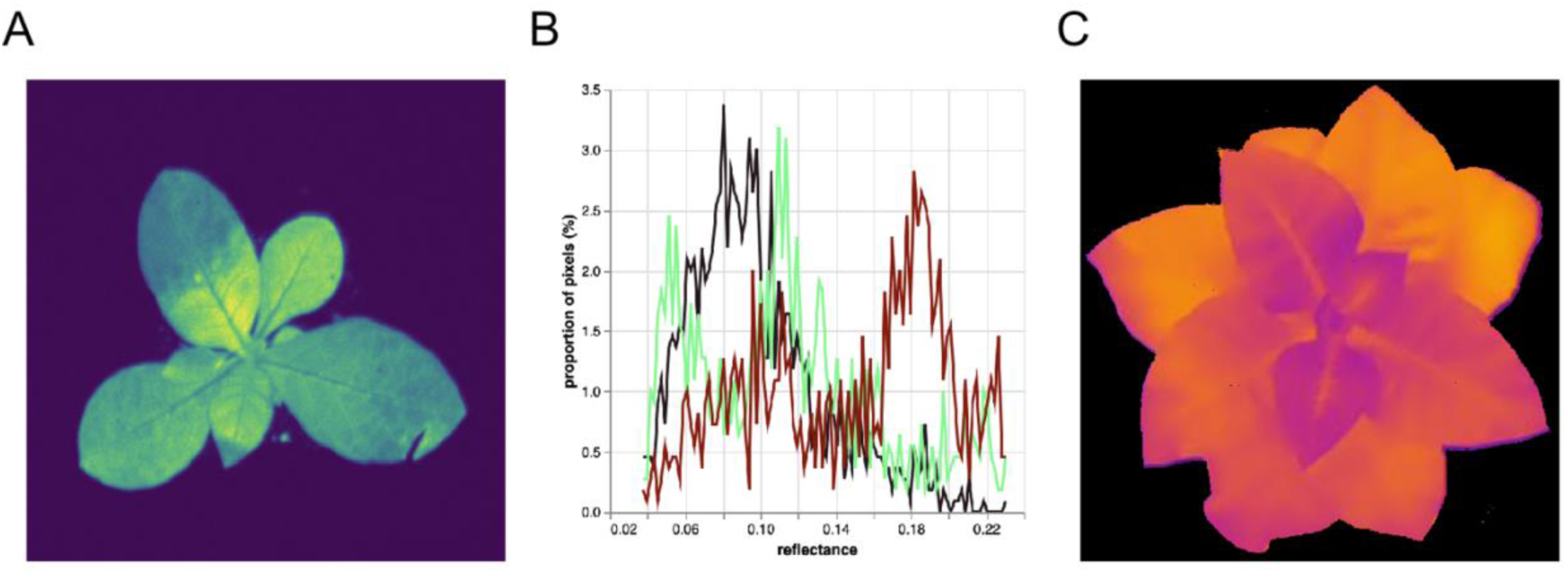
New input image types are supported in PlantCV v4. (A) Fluorescence images, such as this image captured using a Phenovation CropReporter can be used to measure photosynthetic activity. (B) Reading in multi- and hyperspectral images results in storing a datacube, from which histograms of reflectance at multiple wavelengths can be visualized simultaneously. (C) Pseudocolored thermal image after using perspective warp to segment with the accompanying RGB image. Images in (A) and (C) from (Murphy et al., 2025).

#### Hyperspectral imaging and calculated indices

Hyperspectral imaging has been used to estimate plant phenotypes like biomass, yield, stress response, and disease state (reviewed in (Sarić et al., 2022). In PlantCV v4 we have added functions to read and store hyperspectral data, as well as calibrate and calculate a range of spectral indices. Hyperspectral data stored in the environment for visualizing imagery (ENVI) or Esri ASCII formats, which combine an image file and a header file (.hdr file extension) with metadata, can be read and interpreted by PlantCV v4. We created a custom spectral data class that contains information about the datacube including the wavelengths measured, the datatype, interleave type, and filename. The spectral data class is new to PlantCV v4, and functions that use it downstream benefit from keeping everything contained in a single object input for simplicity and automation. Because multi and hyperspectral data are more difficult to visualize than grayscale and color images, PlantCV creates a pseudo-RGB image from the available visible spectrum bands. The pseudo-RGB image is made with the first, middle, and last band if the approximate target wavelengths (710 nm for red, 540 nm for green, and 480 nm for blue) are not available.

One approach to extracting biologically meaningful plant traits from hyperspectral image data involves mathematical combinations of values at particular wavelengths to yield spectral indices known to be predictive of various aspects of plant health (see references in Table S1). First, images are calibrated using a white and dark reference image (Shaikh et al., 2021) to calculate the calibration as follows:

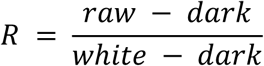

After segmentation of focal objects, users can visualize histograms of particular wavelengths (Figure 6B), calculate predefined spectral indices (Table S1), and output reflectance metrics like mean, median, and standard deviation. These steps function similarly to previously available analysis functions for grayscale and RGB images, but rather than gathering frequencies per pixel value for every band, users can view a histogram based on average reflectance values of each wavelength within a masked area, which is useful for comparing spectra across treatments. Users are not limited to these supported indices; custom spectral indices can be calculated from the user-specified wavelengths. PlantCV searches the available wavelengths to find the closest to the user-requested wavelength within a specified tolerance and stores information about the actual wavelength used for reproducibility.

Incorporating existing hyperspectral image analysis methods into PlantCV is central to maintaining interoperability between PlantCV and other existing open-source phenotyping software. The GatorSense hyperspectral image analysis tools developed by the Machine Learning and Sensing Lab contain a large suite of analysis functions (Zare et al., 2019). Two helper functions were added to PlantCV that calculate average reflectance and the inverse covariance to create an interface between PlantCV hyperspectral tools and GatorSense tools (Zare et al., 2019). Two tutorials on hyperspectral analysis are available (Schuhl et al., 2025b; sdkenney et al., 2025).

#### Thermal imaging

Plant biologists are interested in thermal imaging to measure the temperature of leaves and other organs, which can vary substantially from the air temperature. For example, both abiotic and biotic plant stresses can trigger changes in stomatal closure (Lee and Luan, 2012) leading to changes in leaf temperature due to changes in evaporative cooling via transpiration. PlantCV v4 can directly read temperature values from images taken with Teledyne FLIR thermal cameras. Temperature values, which are estimated from radiometric values and parameters collected by thermal cameras (Sagan et al., 2019), for the focal objects of interest can be analyzed and PlantCV records the mean and median temperatures as well as a histogram of values matching the typical analysis of other data types.

Some cameras collect both thermal and RGB images. Segmentation of plants can be computationally easier using the RGB images, but dual modality cameras like the RGB and thermal sensors often have different resolutions and perspectives such that a labeled mask from the RGB image cannot be applied directly to the thermal image. To enable use of both RGB and thermal data for segmentation, we added perspective warp functionality to PlantCV v4 to align multiple images using a projective transformation (Szeliski, 2010). To align two images, the user provides at least four coordinates of corresponding points (landmarks) in both images, which are used to calculate a projective transformation matrix and to warp one image to match the reference image such that both images are in the same coordinate system. For dual thermal cameras, the RGB image is warped to align it to the reference thermal image. For fixed imaging systems, a transformation matrix can be calculated using stationary landmarks and can be used to transform multiple images without a reference image. Therefore, the transformation matrix just needs to be calculated once per dataset if the camera and target object do not move. A tutorial on thermal image analysis is available (lacostag and Schuhl, 2024).

### 3.4 Additional object segmentation methods

#### 2D Segmentation

A typical workflow for binary segmentation with plant images transforms a RGB image to a colorspace that is perceptually more intuitive (e.g. LAB), then selects a channel that maximizes the difference between plant and background pixels, and finally sets a threshold on the grayscale values where all pixel values are converted to white (included in the mask) or black (excluded from the mask) depending on if the initial pixel value is above or below the threshold value. This approach works well in cases where grayscale values are approximately bimodally distributed (one peak for the background and one for the plant or focal object) (Figure 3C) but is limited because information from only a single color channel can be used. To increase segmentation efficiency by utilizing more image information content, we added a dual channel segmentation tool to PlantCV v4. The dual channel segmentation tool arranges grayscale values from two selected color channels, paired from all supported colorspaces (RGB, LAB, HSV, CMYK), on a 2-dimensional plane. Rather than inputting a threshold value, users input the parameters for a line that bisects the plane and pixels corresponding to points above and below the line are set to black or white to create a mask of the focal objects, functionally segmenting on two color channels simultaneously.

#### K-means clustering

Machine learning (ML) classifiers have been used to successfully separate objects of interest from background in image analysis using color values to classify individual pixels (Lee et al., 2018; Adams et al., 2020; Miao et al., 2020), particularly when single or dual-channel thresholding is insufficient for segmentation. ML classifiers have an added benefit beyond segmentation in that they can provide the number of pixels in the assigned categories, for example diseased vs. healthy plant tissue. PlantCV v2 introduced segmentation using a naive Bayes classifier, a supervised ML model that requires users to train a model based on pixels assigned to each category of interest and has been used successfully in plant phenotyping (Enders et al., 2019; Pierz et al., 2023; Ludwig et al., 2023; Teng et al., 2025). In v4, we have added an unsupervised ML segmentation method using a mini-batch k-means clustering algorithm (Sculley, 2010). Our method applies a Gaussian blur using scikit-image (van der Walt et al., 2014) and extracts neighborhoods around each pixel (termed patches) using scikit-learn (Pedregosa et al., 2011). The mini-batch k-means method is then used to fit a clustering model on the patches, allowing for better inclusion of spatial information within the image.

Our patch-based k-means clustering implementation has two primary parameters: the number of clusters for classification (k) and the size of the patch neighborhood (p). In some circumstances, the combination of k and p can have dramatic effects on the resulting pixel clusters. To make it easier for users to search a subset of parameter space, we have also added a tiling visualization function which produces a composite of segmented images in a grid.

#### Segment image series

A common image analysis challenge is segmentation of individual plants within an image if the plants are touching or overlapping (Gutierrez Ortega et al., 2021). Overlapping plants is especially common in plant growth facilities where plants are grown at high density in trays and overlap as they grow larger, leading traditional segmentation methods to consider multiple plants a single plant for segmentation and downstream analysis. To address this issue, we introduced a watershed segmentation method (Beucher, 1992) that propagates individual plant labels in both space and time dimensions (Gutierrez Ortega et al., 2021). The approach relies on the regularity of timelapse imaging where plants stay in the same place and grow a small amount between timepoints. Starting at the first time point when the plants are small and not overlapping, the initial segmented plants are utilized as markers for watershed segmentation. All images in the time series are stacked and watershed propagates the initial labels spatially within each image, and between successive timepoints. We previously found that this approach was capable of increasing the analyzable size of a dataset by 28% that otherwise could not be analyzed with traditional segmentation (Gutierrez Ortega et al., 2021).

### 3.5 Morphology subpackage for additional measurements of object shape

Leaf angle is a target for breeding and modification in cereal crops because it greatly impacts yield and plant density tolerance (Lambert and Johnson, 1978) but is very time consuming to manually measure. In PlantCV v4, we added a subpackage of modular functions to measure numerous additional morphological traits. These tools are useful to phenotype leaves, stems, hypocotyls, and roots if target objects are not overly complex (i.e. with lots of overlapping and occlusion). With these functions, users can thin a binary mask to a morphological skeleton, identify tips and branching points of those skeletons, break down the skeletons into segments, sort segments, and fill them to create a segmented version of the whole plant or organ (Figure 7A).

**Figure 7.**
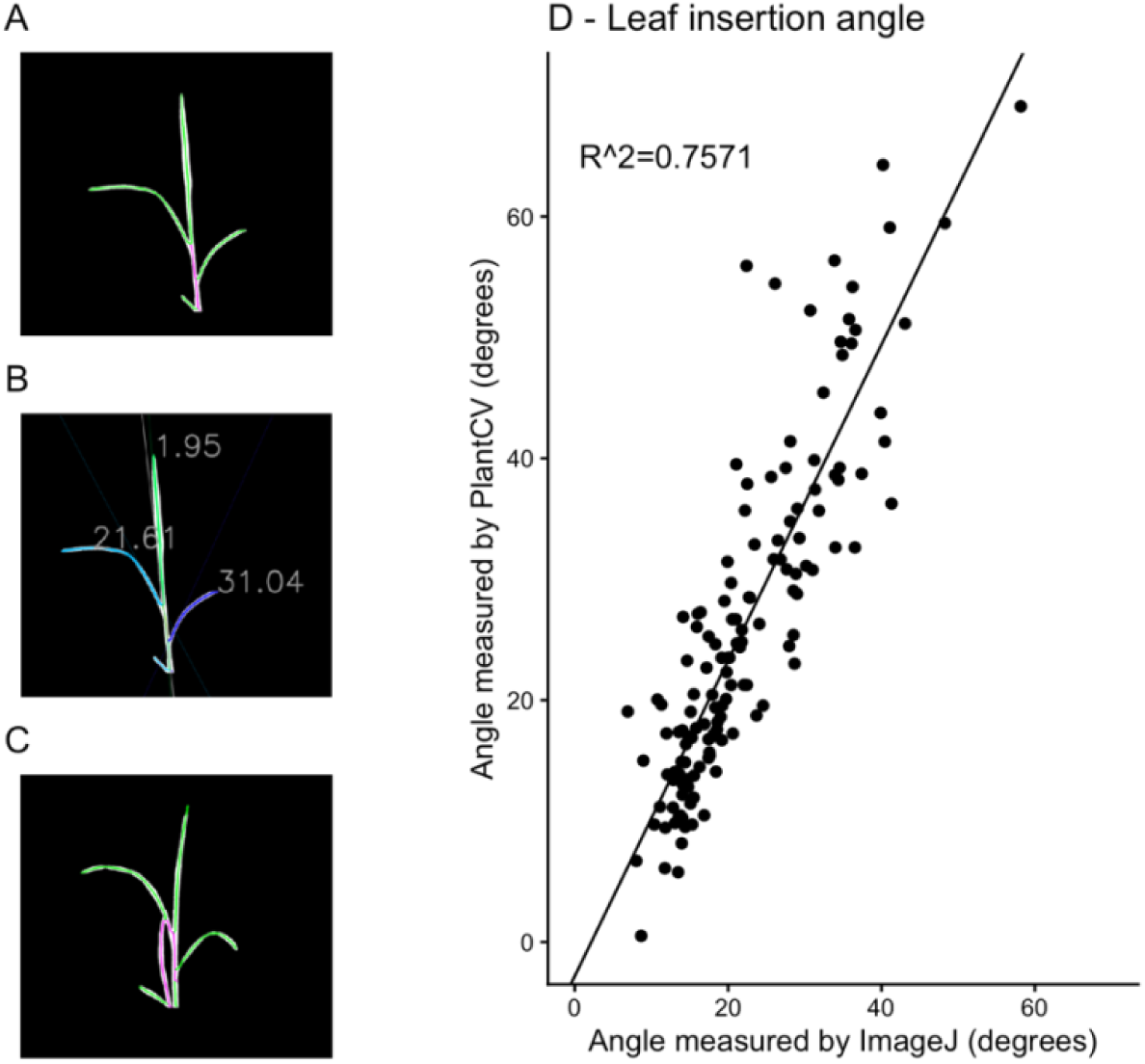
Segment insertion angle measured with PlantCV software. (A) PlantCV finds the central culm and attached leaves of a rice plant from the skeletonized mask. (B) The recorded angle values are calculated by fitting linear regression lines to the inner portion of every leaf segment provided, where the amount of leaf used to fit this linear regression is parameterized by the user as it is dependent on plant species and image resolution. (C) Obscuring or overlapping leaves create closed loops, which causes the segment sorting algorithm to fail since segments that are actually leaf will get categorized as part of the stem. (D) Correlation between PlantCV-measured and ImageJ-measured leaf insertion angle.

We combined the scikit-image skeletonization function (van der Walt et al., 2014) and a plant architecture determination algorithm described to measure leaf angle in maize (Das Choudhury et al., 2018) for segment sorting. However, we maintained the units and positions of segments within the image data array, subtracting branch points to create discrete segments of the plant skeleton, rather than treating the skeleton of the plant as a mathematical graph with nodes and edges, as was done in the original algorithm. The logic is similar, but the PlantCV v4 subpackage uses information about the number of tips and branch points corresponding to each segment to determine whether a segment will be grouped with primary objects (e.g. stems) or secondary objects (e.g. leaves). A tutorial that contains an example workflow that segments and measures leaves is available (Schuhl et al., 2025c).

After skeletonizing and separating segments into primary and secondary objects, several plant phenotypes can be automatically estimated. Length of both stems and leaves is straightforward to calculate from the skeleton, and the angle of leaves can be represented in a variety of ways: angle of insertion relative to the stem, average angle across the whole leaf segment, angle from the leaf inflection point (erectness), and the angle from the leaf tip to the stem. In PlantCV v4, we have made describing the shape of a plant’s skeleton flexible enough to enable estimation of many traits that are time consuming to measure manually but are useful for understanding plant growth. PlantCV morphology tools have also been used for applications outside of the plant science field (Palermo et al., 2021; Kim et al., 2021; Iturburu et al., 2023; Castaño Mansillas, 2024; Sundharbaabu et al., 2024; Rieder et al., 2024).

Leaf segments are ordered based on position within an image rather than a biologically meaningful way (for example, from the bottom of the plant upwards). Ordering by image position does not allow leaf measurements to be tracked over time, or to associate measurements with plant development. We added the ability to automatically resort leaf identifiers for plants imaged from the side view by approximate developmental order (bottom to top; Figure 7A). PlantCV identifies the branch point between each leaf and the stem and orders the branch points along the vertical axis of the image, then re-orders the leaves based on their branch point position from the bottom to the top of the image. This approach will fail if a leaf is missing or undetectable but generally allows for consistent numbering and comparison over time.

The morphology subpackage attempts to measure three-dimensional attributes from two-dimensional representations of data, which has limitations and challenges. Complex structures, such as root systems or plants with multiple tillers or branches, are not amenable to this method (Figure 7B). Further, this approach assumes that the two-dimensional representation of traits like leaf length or angle approximate their true value. For example, leaf length will be accurately calculated when a leaf is parallel to a background surface, but length measurements will be influenced by positioning if a leaf is growing toward or away from the camera.

To understand the impact of these limitations and to examine if the morphology methods described here extract information with biological relevance, we used PlantCV v4 to extract leaf insertion angle with a set of previously published rice images. We compared manually recorded measurements collected with ImageJ against those collected in an automated manner with PlantCV v4. The morphological traits were extracted with a set workflow; human input was used to match the replicate leaves within a plant to their manually measured counterpart for the comparison (Figure 7C). Manually tracing leaves with a software like ImageJ can take 1-5 minutes per image, depending on the plant and number of leaves. Measurements collected with PlantCV closely correlated with manual measurements (R^2^ = 0.7571, t-value of linear regression of PlantCV measurements onto ImageJ measurements = 20.663, p < 2e-16, Figure 7D). One caveat of this rice dataset is that researchers spent time and effort to manually position plants so that it would be at the optimum angle for comparing leaf angle traits.

### 3.6 Interactive documentation and tutorial gallery

In addition to the static documentation hosted with Read the Docs (Gehan et al., 2017), PlantCV v4 now includes interactive documentation with tutorials that allow users to test PlantCV functions and applications without installing any software. Users can launch tutorials in cloud environments using Binder (Jupyter et al., 2018) or Google Colaboratory (Google LLC,), work with included sample data, or upload and analyze their own images by modifying and executing tutorial workflow steps. The ability to use PlantCV without any downloads or installation further lowers barriers in using this software. The PlantCV website (https://plantcv.org) includes a tutorial gallery with use cases, and an updated documentation page on how to contribute a tutorial to the gallery. Examples of user-contributed tutorials include workflows for maize tassel and anther phenotyping (Fahlgren et al., 2025b; Teng et al., 2025), assessing differential growth of *Setaria* leaves using pseudo-landmarks (Hodge et al., 2021; Fahlgren et al., 2025a), and measurement of rice grain chalkiness (Medina Jimenez et al., 2025). Tutorial code and data are hosted in separate GitHub repositories, which allows developers to document the tutorial dependencies for reproducibility. The collection of tutorials includes the majority of all PlantCV functions. Tutorials are versioned and deposited on Zenodo (re3data.org, 2013), and when functions in PlantCV or other dependencies change, the tutorials are updated with new versions for maintenance and sustainability.

Interactive tutorials are also useful for workshops, course-based education, and outreach activities. Each user in a group setting can initiate an independent instance of the exact same Jupyter environment, run the workflow, and add their own changes. Workshops that teach image analysis with PlantCV touch on computer science, mathematics, and biology, and since most workflow steps take less than a second to execute, any updates made to the pre-written code can be quickly visualized and compared. Students can also run the image analysis workflows in the tutorials on their own data, making PlantCV implementation within authentic research experiences accessible. Learning image analysis is inherently visual, and seeing an output image moments after executing a line of code in a Jupyter notebook provides near instant feedback for students, cementing new concepts in real time.

## 4 CONCLUSIONS

PlantCV maintains a common data format that enables interoperability with other image analysis tools, and a modular design that facilitates the rapid assimilation and integration of new methods. PlantCV v4 provides new and updated code for installation, input data types, functions for morphology, numerous tutorials, and accompanying documentation. Open-source software developed in an academic lab is challenging to maintain, as trainees graduate or move to other positions and dependent packages change. The continuous development and maintenance, thanks to a global community of contributors, have enabled the bug fixes, updated backend, new functions, and new tutorials described here. As new tools in Deep Learning come online, researchers must be able to validate the tools and make meaningful biological conclusions, maintaining the relevance of PlantCV. Continuing to connect PlantCV to other interoperable, maintained tools, as well as build out new functionality for plant phenotyping, will enable further progress in PlantCV and the field of open-source phenotyping software.

## ABBREVIATIONS

BIL: Band Interleaved by Line
BIP: Band Interleaved by Pixel
BSQ: Band Sequential
CMYK: Cyan Magenta Yellow Black
ENVI: Environment for Visualizing Imagery
HSV: Hue Saturation Value
JSON: JavaScript Object Notation
LAB: Lightness, Red/Green, Blue/Yellow
MIAPPE: Minimum Information About a Plant Phenotyping Experiment
ML: Machine learning
NIR: Near Infrared
RGB: Red Green Blue
ROI: Region of interest

## ACKNOWLEDGMENTS

We would like to thank the PlantCV users and community. This work is supported by the Donald Danforth Plant Science Center, USDA-NIFA (2019-67021-29926 and 2022-67021-36467 to N.F. and M.G.; 2022-67013-36129 and 2021-67013-33778 to M.G.; 2020-67034-31901 to A.C.), DOE (DE-SC0018072 to N.F and M.G. and DE-SC0023142 to M.G.), NSF (2121293, 2120153, 2346101, 2347188, and 1921724 to N.F. and M.G.), the Taylor Geospatial Institute, and the Wells Fargo IN^2^ program. This work was supported by Defense Advanced Research Projects Agency Advanced Plant Technologies (DARPA-APT, HR001118C01327). We thank the Plant Growth Facility (RRID:SCR_024902) and the Phenotyping Core Facility (RRID:SCR_019049) at the Donald Danforth Plant Science Center for plant growth and image collection, respectively. The views, opinions, and /or findings expressed are those of the authors and should not be interpreted as representing the official views or policies of the Department of Defense of the U.S. Governments. A.M. has recently joined the European Research Council Executive Agency (ERCEA) as a scientific officer. The information and views set out in this article are those of the author and do not necessarily reflect the official opinion of the European Commission and ERCEA).

## DATA AVAILABILITY STATEMENT

Links to code, tutorials, documentation, and other resources are available on the PlantCV homepage at https://plantcv.org. PlantCV source code is available on GitHub at https://github.com/danforthcenter/plantcv. Scripts used for analyses in this paper are available on GitHub at https://github.com/danforthcenter/plantcv-4-paper.

