## Supplemental Materials for "PlantCV v4: Image analysis software for high-throughput plant phenotyping"

**Table S1**. **Supported spectral indices in PlantCV**. In the equations, "Rxx" represents reflectance at a specific wavelength band. In PlantCV, reflectance at 800nm (R800) is used as near infrared (NIR) reflectance; reflectance at 670 (R670) is used as RED reflectance; reflectance at 550 (R550) is used as GREEN reflectance; reflectance at 480 (R480) is used as BLUE reflectance. Specifically for EGI, r, g, and b represents for ratios for red, green and blue, respectively, i.e. r = R / (R + G + B), g = G / (R + G + B), b = B / (R + G + B).

| Spectral Index | Abbreviation | Formula | Reference | Value Range |
|---|---|---|---|---|
| Anthocyanin Reflectance Index | ARI | (1/R550)-(1/R700) | (Gitelson et al., 2001) | (-∞, ∞) |
| Chlorophyll Index Red-edge | CI_red-edge | (R800/R700)-1 | (Gitelson et al., 2003) | (-1, ∞) |
| Carotenoid Reflectance Index 550 | CRI550 | (1/R510)-(1/R550) | (Gitelson et al., 2002b) | (-∞, ∞) |
| Carotenoid Reflectance Index 700 | CRI700 | (1/R510) - (1/R700) | (Gitelson et al., 2002a) | (-∞, ∞) |
| Excess Green Index | EGI | 2g - r - b | (D. M. Woebbecke et al., 1995) | (-1.0, 2.0) |
| Enhanced Vegetation index | EVI | (2.5 * (NIR - RED)) / (1 + NIR + (6 * RED) - (7.5 * BLUE)) | (Huete et al., 1997) | (-∞, ∞) |
| Green Difference Vegetation Index | GDVI | (NIR - GREEN) / (NIR + GREEN) | (Sripada et al., 2006) | (-2.0, 2.0) |

| | | | | |
|---|---|---|---|---|
| Modified Anthocyanin Reflectance Index | MARI | ((1 / R550) - (1 / R700)) * R800 | (Gitelson et al., 2006) | (-∞, ∞) |
| Modified Chlorophyll Absorption Reflectance Index | MCARI | ((R700 - R670) - 0.2 * (R700 - R550)) * (R700 / R670) | (Daughtry et al., 2000) | (-∞, ∞) |
| MERIS Terrestrial Chlorophyll Index | MTCI | (R753.75 - R708.75) / (R708.75 - R681.25) | (Dash and Curran, 2004) | (-∞, ∞) |
| Normalized Difference Red Edge Index | NDRE | (R790 - R720) / (R790 + R720) | (Barnes et al., 2000) | (-1.0, 1.0) |
| Normalized Difference Vegetation Index | NDVI | (NIR - RED) / (NIR + RED) | (Rouse et al., 1974) | (-1.0, 1.0) |
| Photochemical Reflectance Index | PRI | (R531 - R570) / (R531 + R570) | (Penuelas et al., 1995b) | (-1.0, 1.0) |
| Pigment Specific Normalized Difference for Chlorophyll a | PSND_CHLa | (R800 - R680) / (R800 + R680) | (Blackburn, 1998) | (-1.0, 1.0) |
| Pigment Specific Normalized Difference for Chlorophyll b | PSND_CHLb | (R800 - R635) / (R800 + R635) | (Blackburn, 1998) | (-1.0, 1.0) |
| Pigment Specific Normalized Difference for Caroteniods | PSND_CAR | (R800 - R470) / (R800 + R470) | (Blackburn, 1998) | (-1.0, 1.0) |
| Plant Senescence Reflectance Index | PSRI | (R678 - R500) / R750 | (Merzlyak et al., 1999) | (-∞, ∞) |

| | | | | |
|---|---|---|---|---|
| Pigment Specific Simple Ratio for Chlorophyll a | PSSR_CHLa | R800 / R680 | (Blackburn, 1998) | (-1.0, 1.0) |
| Pigment Specific Simple Ratio for Chlorophyll b | PSSR_CHLb | R800 / R635 | (Blackburn, 1998) | (-1.0, 1.0) |
| Pigment Specific Simple Ratio for Carotenoids | PSSR_CAR | R800 / R470 | (Blackburn, 1998) | (-1.0, 1.0) |
| Red:Green Ratio Index | RGRI | RED / GREEN | (Gamon and Surfus, 1999) | (0.0, ∞) |
| Red-Edge Vegetation Stress Index | RVSI | ((R714 + R752) / 2) - R733 | (Merton and Huntington, 1999) | (-1.0, 1.0) |
| Soil Adjusted Vegetation Index | SAVI | (1.5 * (NIR - RED)) / (NIR + RED + 0.5) | (Huete, 1988) | (-1.2, 1.2) |
| Structure-Independent Pigment Index | SIPI | (NIR - RED) / (NIR - BLUE) | (Penuelas et al., 1995a) | (-∞, ∞) |
| Simple Ratio | SR | NIR / RED | (Jordan, 1969) | (0.0, ∞) |
| Visible Atmospherically Resistant Index | VARI | (GREEN - RED) / (GREEN + RED - BLUE) | (Gitelson et al., 2002a) | (-∞, ∞) |
| Vegetation Index using green bands | VI_green | (GREEN - RED) / (GREEN + RED) | (Gitelson et al., 2002a) | (-1.0, 1.0) |
| Water Index | WI | R900 / R970 | (Penuelas et al., 1997) | (0.0, ∞) |

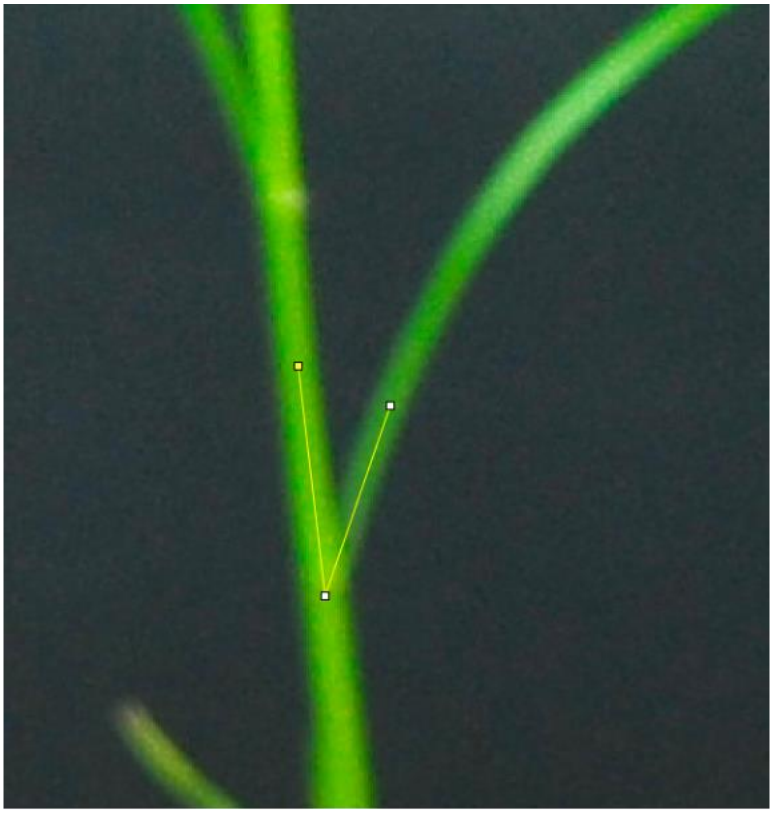

**Figure S1**. **Leaf insertion angle as measured by ImageJ**.
